# Human monocyte subtype expression of neuroinflammation and regeneration-related genes is linked to age and sex

**DOI:** 10.1101/2024.03.10.584323

**Authors:** Juliane F. Tampé, Emanuela Monni, Sara Palma-Tortosa, Emil Brogårdh, Charlotta Böiers, Arne G. Lindgren, Zaal Kokaia

**Affiliations:** Laboratory of Stem Cells and Restorative Neurology, Lund Stem Cell Center, Lund University, Lund, Sweden; Division of Molecular Hematology, Lund Stem Cell Center, Lund University, Lund, Sweden; Department of Clinical Sciences Lund, Neurology, Skåne University Hospital, Lund University, Lund, Sweden

## Abstract

Stroke is a leading cause of disability and the third cause of death. The immune system plays an essential role in post-stroke recovery. After an ischemic stroke, monocytes infiltrate the injured brain tissue and can exacerbate or mitigate the damage. Ischemic stroke is more prevalent in the aged population, and the aging brain exhibits an altered immune response. There are also sex disparities in ischemic stroke incidence, outcomes, and recovery, and these differences may be hormone-driven and determined by genetic and epigenetic factors. Here, we studied whether human peripheral blood monocyte subtype (classical, intermediate, and non-classical) expression of neuronal inflammation- and regeneration-related genes depends on age and sex. A FACS analysis of blood samples from 44 volunteers (male and female, aged 28 to 98) showed that in contrast to other immune cells, the proportion of natural killer cells increased in females. The proportion of B-cells decreased in both sexes with age, and subtypes of monocytes were not linked to age or sex. Gene expression analysis by qPCR identified several genes differentially correlating with age and sex within different monocyte subtypes. Interestingly, *ANXA1* and *CD36* showed a consistent increase with aging in all monocytes, specifically in intermediate (*CD36*) and intermediate and non-classical (*ANXA1*) subtypes. Other genes (*IL-1β, S100A8, TNFα, CD64, CD33, TGFβ1, TLR8, CD91*) were differentially changed in monocyte subtypes with increased aging. Most age-dependent gene changes were differentially expressed in female monocytes. Our data shed light on the nuanced interplay of age and sex in shaping the expression of inflammation- and regeneration-related genes within distinct monocyte subtypes. Understanding these dynamics could pave the way for targeted interventions and personalized approaches in post-stroke care, particularly for the aging population and individuals of different sexes.

## INTRODUCTION

The brain’s innate immune system is mainly comprised of microglia. These specialized cells act as the first line of defense against invading pathogens and play a critical role in maintaining normal brain function [1].

Monocytes are a heterogeneous population [2] with pro- or anti-inflammatory phenotypes depending on the stage of differentiation and mechanism by which they are activated [3]. Under normal physiological conditions, the blood-brain barrier (BBB) generally prevents blood-circulating monocytes and other immune cells from entering the brain tissue. Brain damage such as traumatic injury or stroke opens the BBB, and immune cells from the blood, including monocytes, may enter the brain tissue [4]. After infiltration into the stroke-damaged brain, the blood-circulating monocytes are recruited to the site of the ischemic brain injury, become macrophages, and, together with activated microglia, contribute to neuroinflammation and impairment/regeneration of brain function [5–7]. Studies have shown that monocyte-derived macrophages (MDM) can influence inflammation and tissue damage by releasing pro-inflammatory cytokines such as tumor necrosis factor-alpha (*TNF-α*), interleukin-1β (*IL-1β*), and matrix metalloproteinase-9 (*MMP9*) [8, 9]. These cytokines contribute to activating immune cells, oxidative stress, and neuronal damage [10]. MDMs can also have anti-inflammatory phenotypes and promote brain tissue and functional regeneration by releasing factors such as *TGF-β*, *BDNF*, and *IL10* [1, 4].

We previously showed in an animal model [11] that depletion of MDM during the first week after stroke abolished long-term behavioral recovery and drastically decreased tissue expression of anti-inflammatory genes, including *TGF-β*, *CD163*, and *Ym1*. Our observations suggest that MDMs play an essential role in post-stroke recovery by activating anti-inflammatory factors. In support, we later reported potentially clinically important data showing that mouse MDMs, primed *in vitro* to become anti-inflammatory macrophages and then administered into cerebrospinal fluid of stroke-subjected mice, infiltrate into the ischemic hemisphere and promote post-stroke recovery of motor and cognitive functions [12].

In contrast to mice, human monocytes are identified by expressing specific cell surface markers, such as *CD14* and *CD16. CD14* is a glycoprotein that recognizes bacterial lipopolysaccharides and is present in most monocytes, whereas *CD16* is a low-affinity Fcγ receptor. Based on the expression of these surface markers and functional characteristics, human monocytes are commonly categorized [13] into the three subtypes. Classical monocytes (CD14+/CD16−) are the most abundant subtype in circulation, representing about 85% of the total number of monocytes [14]. They are characterized by the high expression of *CD14* and the low expression of *CD16* [15]. These classical monocytes are known for their phagocytic and antigen-presenting capabilities. Non-classical monocytes (CD14−/CD16+) comprise about 10% of total monocytes. They are characterized by the low expression of *CD14* and the high expression of *CD16*. Non-classical monocytes are known for their patrolling functions, surveillance of endothelial surfaces, and their role in the early immune response to infections [16]. Intermediate monocytes (CD14+/CD16+) are relatively rare, comprising about 5% of the total monocytes [17, 18]. Intermediate levels of *CD14* and *CD16* expression are characteristic of this subtype of monocytes, and they are potent producers of inflammatory cytokines contributing to acute and chronic inflammation [19].

To keep these different functions in steady-state homeostasis throughout the whole body, these three monocytes’ subsets are continuously produced and circulate through the bloodstream in a dynamic equilibrium: human monocytes arise within 1-2 days from the bone marrow as classical monocytes for one day. This process depends on the expression of the chemokine receptor *CCR2* [20, 21] and is specific to classical monocytes [22]. Afterwards, they transition for 4 days as intermediate monocytes towards non-classical monocytes, which survive for 7 days [23]. This process results in a constant pool of 85% classical, 5% intermediate, and 10% non-classical monocytes [14, 24, 25]. *CD91* has been identified as a marker for all monocytes and is steadily expressed through all subtype stages of the differentiation [26].

Similar to mice [27], acute brain damage, such as stroke, spinal cord injury, and traumatic brain injury (TBI), significantly impacts the peripheral immune system in humans [28–30]. Increased numbers of circulating monocytes have been reported in stroke and TBI patients [31, 32] but not in patients with spinal cord injury [33]. Peripheral blood monocytes before and at different time points after a stroke could contribute to a patient’s regenerative potential. Different subtypes of monocytes have been correlated with outcome severity in patients following stroke [34]. However, these monocyte subsets’ pro- or anti-inflammatory functions depend on the inflammatory context, and there is no strict association between phenotype and function [35, 36].

It is still not fully understood how the age and sex of patients can influence the expression of specific genes linked to the inflammation-caused exaggeration of ischemic damage and promotion of regeneration and functional recovery [37, 38]. Monocytes, like most cells in the human body, do not have a defined age. Still, monocytes have a limited life span once differentiated from hematopoietic progenitor cells in the bone marrow [16]. It is still unclear whether the phenotypic and genetic profiles of peripheral monocytes, like most cells in the human body, are related to the monocyte age. In addition, their numbers and function may be affected by the age and sex of the patient. The number of monocytes in circulation may decrease with age [39], and the function of monocytes may also decline, impairing the immune response and increasing susceptibility to infections [40]. Sex may also play a role in the number and function of monocytes [41]. Studies have found that women generally have higher numbers of circulating monocytes than men, but the function of monocytes may be more robust in men than in women [42]. Monocyte counts may also be related to ethnicity. Overall, the effects of age and sex on monocytes are complex and can be influenced by various factors, such as genetics, lifestyle, and environmental exposures [43]. It is unclear whether the age and gender of patients can influence the expression of specific genes linked to the inflammation-caused exaggeration of ischemic damage and promotion of regeneration and functional recovery. Further research is needed to fully understand the properties of monocytes from age and sex perspectives [44].

Here, we studied whether human peripheral blood monocyte subtype expression of inflammation- and regeneration-related genes depends on age and sex. Gene expression analysis identified several genes differentially correlating with age and sex within different monocyte subtypes.

## MATERIALS AND METHODS

### Blood sample collection

The blood samples were collected from male and female volunteers aged between 28 and 98 years (Supplementary Table 1). Pregnancy, systemic inflammatory diseases (e.g., rheumatoid arthritis, systemic lupus erythematosus), or hematologic diseases affecting the immunological status served as exclusion criteria. In case of exclusion from the study, already collected samples were destroyed and discarded.

All procedures were carried out according to the ethical permit obtained from the Local Ethical Committee (ethical permit Dnr 2016/179; 2017/357; and 2017/879) in compliance with the Declaration of Helsinki. This cohort study includes no intervention except clinical examinations and venous blood sampling, carried out using standardized, routine methods. Specialized research nurses at the Department of Neurology at Skåne University Hospital collected 10mL of venous blood in heparin-treated, ethylenediaminetetraacetic acid (EDTA) coated vacutainers (Sarstedt, Germany). All samples were coded not to carry any personal identifier information. To avoid potential bias, samples were randomized, and data collection and analysis were performed blindly.

### Preparation of peripheral blood mononuclear cells

After collection, the blood samples were kept at room temperature (RT) before mononuclear cell isolation within 24h. Peripheral blood mononuclear cells (PBMCs) were isolated in a biosafety level 2+ cell laboratory under constant airflow in a laminar flow hood. All reagents used were devoid of animal-derived products. Human recombinant albumin (HAS) was used in place of fetal bovine serum to avoid any trigger of monocyte activation. All procedures were performed at RT. According to the manufacturer’s instructions, PBMCs were isolated using SepMate tubes (StemCell Technologies, UK) containing Lymphoprep (Serumwerk, Germany) by density gradient centrifugation. PBMCs were frozen in StemCellBanker (Amsbio, UK) at −80 °C. The next day, samples were cryopreserved in liquid nitrogen (−170°C) or −155°C freezer for long-term storage until further analysis.

### Flow cytometry and FACS preparation

The frozen PBMCs were thawed at RT and washed twice in 10 mL buffer. Dulbecco’s Phosphate-Buffered Saline (DPBS) with 2% HSA was used as a buffer during all procedures. The pelleted cells were then resuspended in 60 μL buffer and incubated with the appropriate antibody concentration on a shaker for 30-60 min at 4°C. Then, cells were washed twice with 1 mL of buffer and centrifuged at 500 x g for 5 min. Finally, the resulting pellet was resuspended in 200 μL buffer and strained through the 35 µm mesh incorporated in a tube for flow cytometry applications (Falcon, Corning, USA).

Primary human monocytes and their subtypes were identified and isolated from PBMCs based on their expression of *CD91* [26] and their differential expression of the *CD14* and *CD16* cell surface markers. The B-natural killer (NK)- and T-cells were identified based on their *CD19*, *CD56*, and *CD3* expression. The following antibodies were employed: CD91-PE (clone A2MR-α2), CD14-APC (clone M5E2), CD16-BV421 (clone 3G8), CD3-PE-Cy7 (clone UCHT1), CD19-BB515 (clone HIB19), CD19-PE-Cy7 (clone SG25C1), CD56-BV605 (clone B159), (BD Biosciences, Sweden). To exclude non-viable cells, DRAQ7 (BD Biosciences, Sweden) was added to the cell suspension 15 minutes before analysis (Supplementary Table 2). Cell type identification and isolation were performed using a BD FACSAria™ II cell sorting system (BD Biosciences, Sweden).

To define a gating strategy, unstained cells served as a negative control, and single-stained samples were used to compensate for the fluorophores’ spectral overlap. All cell population types have additionally been confirmed by Fluorescence Minus One (FMOs), re-analysis, and back gaiting. The antibodies and staining volumes have been scaled according to the number of PBMCs to achieve the same staining concentration in all samples (Supplementary Table 3).

### FACS analysis and sorting

The cells were separated from debris and selected for size using the area of the forward (FSC) and side scatter (SCC). We excluded doublets by using FSC-W/H and SCC-W/H, and eliminated dead cells by utilizing the intracellular dye DRAQ7. The three monocyte subpopulations were further classified by the expression of *CD14* and *CD16*: CD14+ and CD16− classical monocytes, CD14-and CD16+ non-classical monocytes, and CD14+ and CD16+ intermediate monocytes. For monocytes and each of the three subtypes, biological duplicates of 20 cells from the same donor were sorted into each well of a 96-well-cell culture plate (Corning, Sweden). Before cell sorting, a lysis buffer consisting of 10% NP40 (Thermo Fisher, Sweden), 10mM dNTP (Takara, Japan), 0.1M DTT (Thermo Fisher Scientific, Sweden), RNaseOUT (Thermo Fisher Scientific, Sweden), and nuclease-free water was dispensed to a 96-well-cell culture plate. The plate was spun at 1300 rpm and kept at −80°C for subsequent pre-amplification and analysis.

### High-throughput microfluidics technology quantitative PCR (Fluidigm)

To analyze the potential age- and sex-related differences in gene expression between the monocytes and their subtypes, we performed a Fluidigm-based study and examined the expression of 39 brain inflammation- and regeneration-related genes. Fluidigm is a microfluidics technology that allows simultaneous and efficient gene expression analysis using integrated fluidic circuits (IFCs) [45]. The IFC chosen performs precise and high-throughput quantification of mRNA levels of up to 96 genes. The small format reduced sample and reagent requirements, allowing sensitive measurements of low mRNA samples. The chosen regenerative genes were selected through an extensive literature review, focusing on genes expressed by monocytes and/or macrophages and implicated in neuroinflammation and the context of stroke recovery. Three housekeeping genes, four negative controls for the other PBMC populations, and one technical control were deployed to normalize the expression and do quality control (Supplementary Table 6).

The complementary DNA (cDNA) was generated with each of the 47 TaqMan probes (S1 Table 5) and Xeno primer (TaqMan Cells-to-Ct Control kit, Thermo Fisher Scientific, Sweden) and using the Taq-SSIII reaction mix (CellsDirectTM One-Step qRT-PCR Kit, Ambion, Thermo Fisher Scientific, Sweden). The negative, positive, no reverse transcriptase (noRT), and linearity controls were included on two plates and only repeated when a new batch of probes or SSIII enzyme was used. We used a polymerase chain reaction (PCR) starting with an extended 50°C for 1 hour followed by 2 minutes at 95°C. For the panel of primers used, we identified that 18 cycles at 95°C for 15 seconds and 60°C for 4 minutes were optimal for pre-amplification. The produced cDNA was used directly for rtPCR or stored at −80°C.

The TaqMan Gene Expression Master Mix (Thermo Fisher Scientific, Sweden) and GE Sample loading reagent (100-7610, Fluidigm, Standard Biotools Inc., California) were mixed 10:1, and 3.3 μL of the mix was added to each well of a 96-well plate. Next, the cDNA from the pre-amplification was diluted 1:5 and 2.7 μL added to the pre-mixed Sample-plate for a final volume of 6 μL. Once all sample and assay mixes were prepared, the Integrated Fluidic Circuit (IFC) chip was injected with the control line fluid via the two valves on the chip. The IFC chip was then inserted into the Integrated Fluidic Unit (IFU) and primed using pressure to push the control line fluid into all reaction chambers and channels, ensuring they were free from air bubbles and debris. Once primed, the chip was loaded by pipetting at least 4 μL of either sample or assay mix into the corresponding wells of the chip. Qualitative gene expression was measured by detecting FAM-MGB during a cycle of 50-70-25-50-96. We set a baseline for the lowest and threshold fluorescence signals using the BioMark HD Data Analysis software (Standard Biotools Inc., California). The resulting cycle threshold (CT) values were normalized against the beta-actin (*ACTβ*) reference gene. The Fluidigm data were used to conduct the linear regression analysis and identify potential correlations between age and the expression of selected genes linked to the monocytes’ function or activation mode and their subtypes or their inflammatory response during brain injury.

### Statistical analysis

In this exploratory study, PBMC samples were block-randomized and blinded before any analysis, so no biases are to be declared. One million events were collected per sample to estimate the overall cell population via FACS. The group comparison between the Adult and Older group (above 65 years), or male and female, was assessed by an unpaired t-test. The rtPCR data of each Fluidigm IFC was analyzed using the BioMark HD analysis software with a quality threshold of 0.65, linear baseline correction, and automatic global cycle threshold. The data from all chips, metadata, and FACS data were pooled using the software R (Supplementary Data). Non-detected runs were set as CT value = 35, (McCall, McMurray et al. 2014), and failed runs were multiplied imputed using Gene, Subtype, Sex, and Age [46]. We normalized both technical and biological duplicates using ACTβ expression and calculated the mean relative expression (rE) from these four values. The association of age with gene expression was assessed using Pearson’s correlation. Residual analysis identified outliers, defined as values more than 3 times the standard deviation, which were excluded from the final analysis. The Pearson correlation coefficient (r), displayed in the heatmap, quantifies the strength and direction of this relationship. A p-value < 0.05 was considered statistically significant. For significant age associations, a fitted linear regression was plotted with a 95% confidence interval, describing the true mean of the linear correlation with 95% certainty.

### Graphics and Design

BioRender.com was used to create Fig. 1., and PRISM (Version 9, GraphPad) was used to plot PBMC cell populations in Fig 2. The heatmap in Figure 6, showing the Pearson correlation coefficient, was generated using the matrix visualization and analysis software Morpheus (https://software.broadinstitute.org/morpheus). All other figures and tables were generated using the software R (R Core Team (2021). R: A language and environment for statistical##computing. R Foundation for Statistical Computing, Vienna, Austria. ##URL https://www.R-project.org/.).

**Fig 1.**
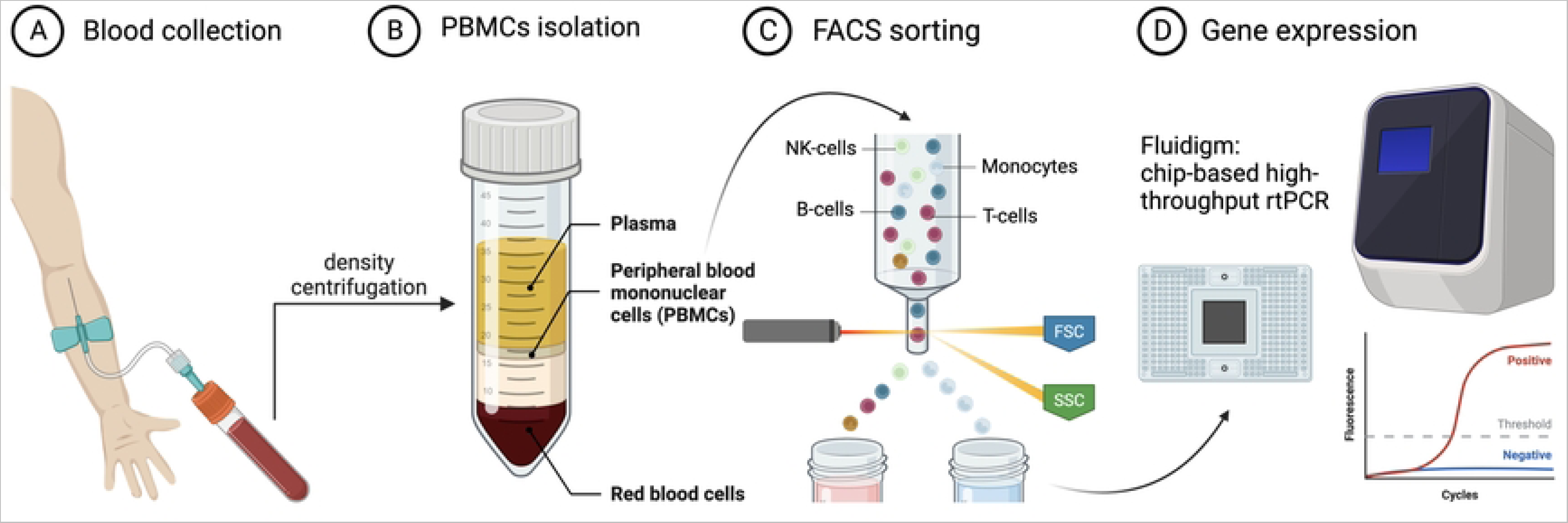
Workflow of the study. (**A**) Blood sample collection. (**B**) Isolation of peripheral blood mononuclear cell isolation (**C**) FACS analysis and isolation of monocytes. (**D**) Gene expression analysis using Fluidigm - a multiplex qPCR. This image was generated using BioRender.com

**Fig 2.**
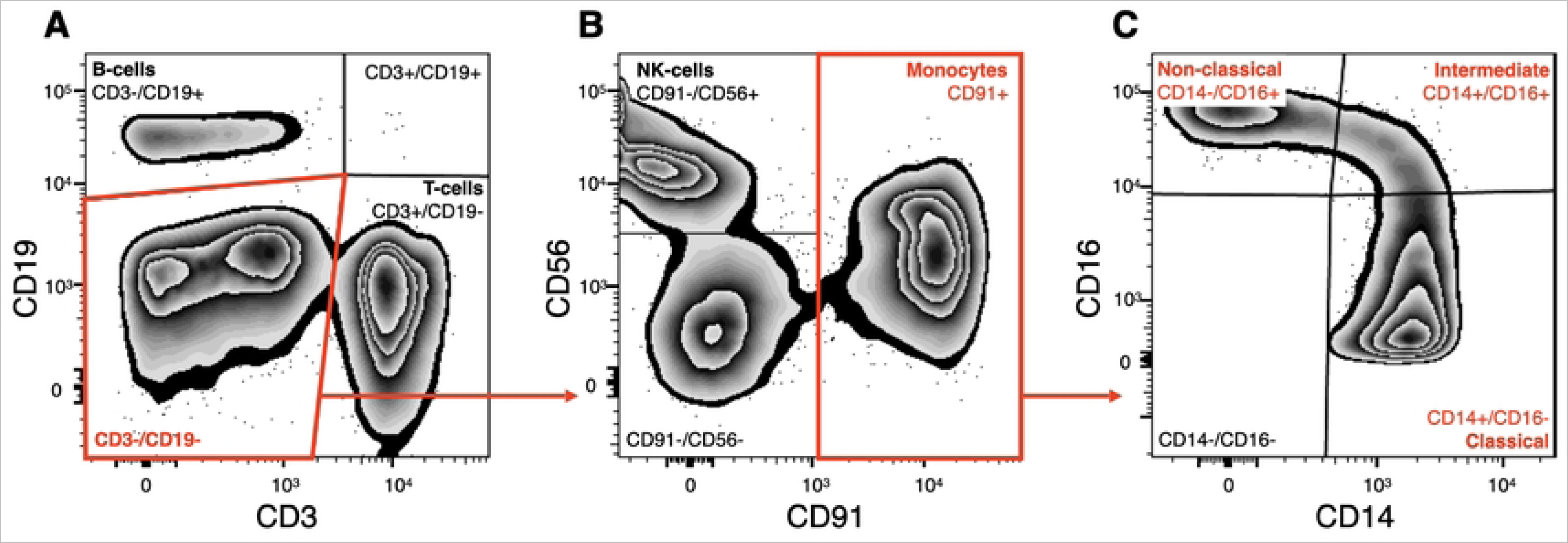
FACS gating strategy of isolated PBMCs and sorting and analysis of the three monocyte subtypes. Single, (Draq7-) cells were considered viable. Cells negative for (**A**) T-cell marker *CD3*, B-cell marker *CD19,* were further gated (**B**) for CD56 (NK-cells) and CD91 (monocytes) (**C**) The CD91+ cells where further sub-fractioned based into classical (CD14+/CD16−), intermediate (CD14+/CD16+), and non-classical (CD14−/CD16+) monocytes.

## RESULTS

The study’s main flow and different steps are presented in Fig 1. The blood samples were obtained from a total of 44 volunteers (18 females (41%) and 26 males (59%) (Supplementary Table 1). The median age for the females was 72 years (range: 43 to 98), while the median age for the males was 66 (range of 28 to 87 years). There was no age variance between male (n=26) and female (n=18) groups (Unpaired t-test; p=0.0634).

### The ratios of B- and NK-cells are linked to age and sex

After isolating PBMCs, we performed a FACS analysis to compare the relative ratio of different PBMCs expressed as a proportion of all PBMCs in the age- and sex-dependent groups. First, we analyzed the expression of *CD19* and *CD3* in PBMCs (Fig 2 A), which allowed us to quantify the ratios of B- (CD3−/CD19+) and T- (CD3+/CD19−) cells. Then, we further analyzed CD3-/CD19- cells for the expression of *CD56* and *CD91* (Fig 2 B) and quantified the proportion of NK-cells (CD56+/CD91−) and monocytes (CD91+). Finally, from the monocyte population, we analyzed subtypes of monocytes based on *CD16* and *CD14* expression (Fig 2 C).

When analyzing the data separated by the donors’ sex, we detected a significant correlation between the female population’s age and the NK-cell ratio (P = 0.025). Additionally, we revealed a significant age-related decrease in B-cells within the viable female (P = 0.016) and male (P=0.013) PBMCs (Fig 4, Supplementary Table 7).

We found no differences in PBMC population proportions when comparing the Adult and Older groups. Adult donors had 11.1±4.8 % B-cells, 21.6±9.8 % NK-cells, 38.9± 8.2 % T-cells, and 18.9± 6.8 % monocytes. In the aged population, we found similar ratios: 10.9±14.6 % B-cells, 29.3±15.2 % NK-cells, 38.0±16.2 % T-cells, and 19.9±10.6 % monocytes (S1 Table 5, S2 Figure 1). When using Pearson’s linear regression method to correlate ratios of the different PBMCs with age, we found that while the total viable PBMCs (P = 0.0003) and B-cells (P=0.003) decreased, the NK-cells (P = 0.043) increased (Fig 3, Supplementary Table 7).

**Fig 3.**
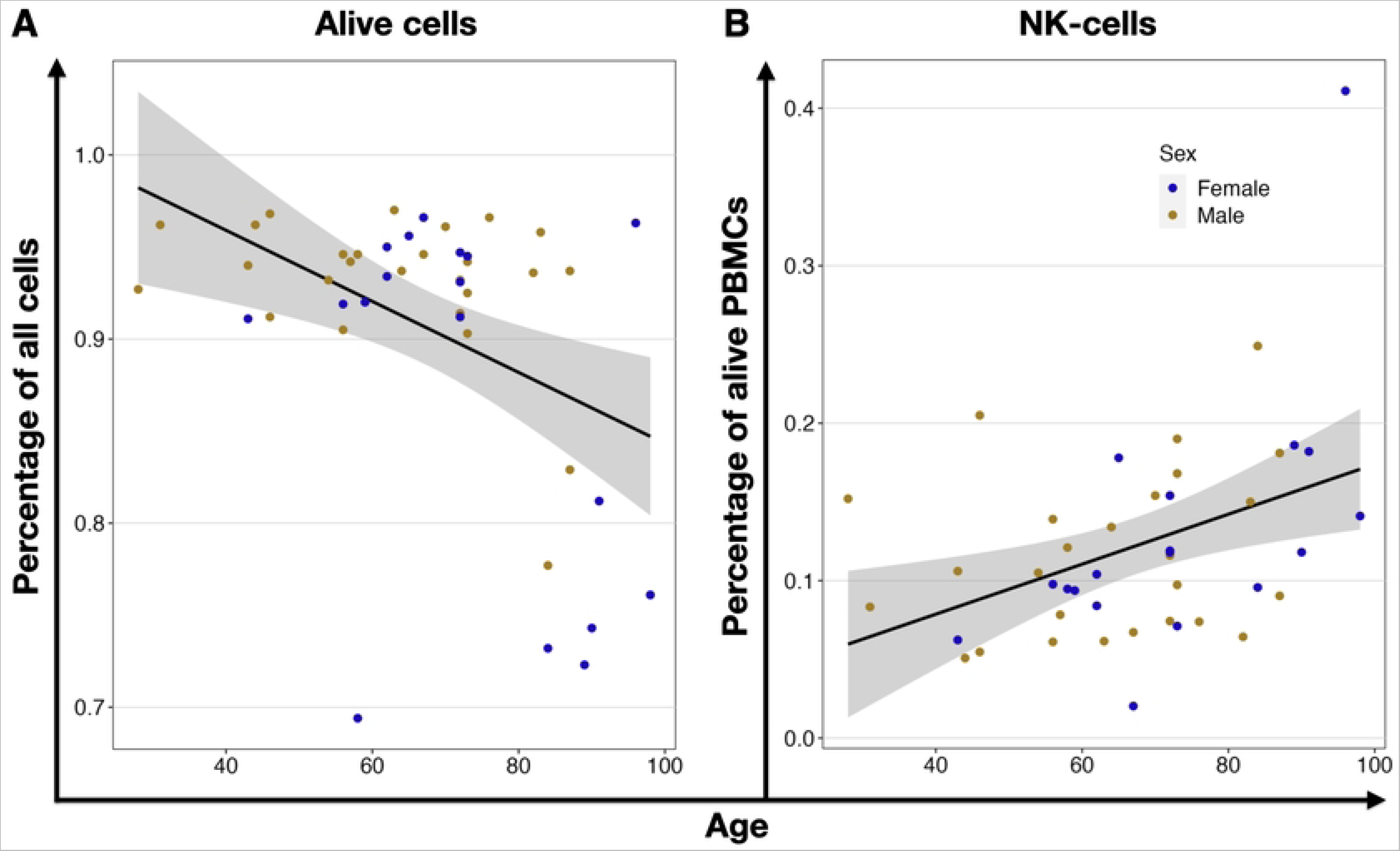
Age-related correlation of PBMCs ratios. (**A**) Shows the decreased (r=−0.52) ratio of Draq7 negative, viable cells in isolated PBMCs, (**B**) decrease (r=−0.44) of B-cells ratio, and (**C**) an increased (r=0.31) ratio of NK-cells within viable PBMCs. Each dot represents an individual donor, yellow for males and blue for females. The black line shows linear regression, and the grey area is the 95% confidence interval.

**Fig 4.**
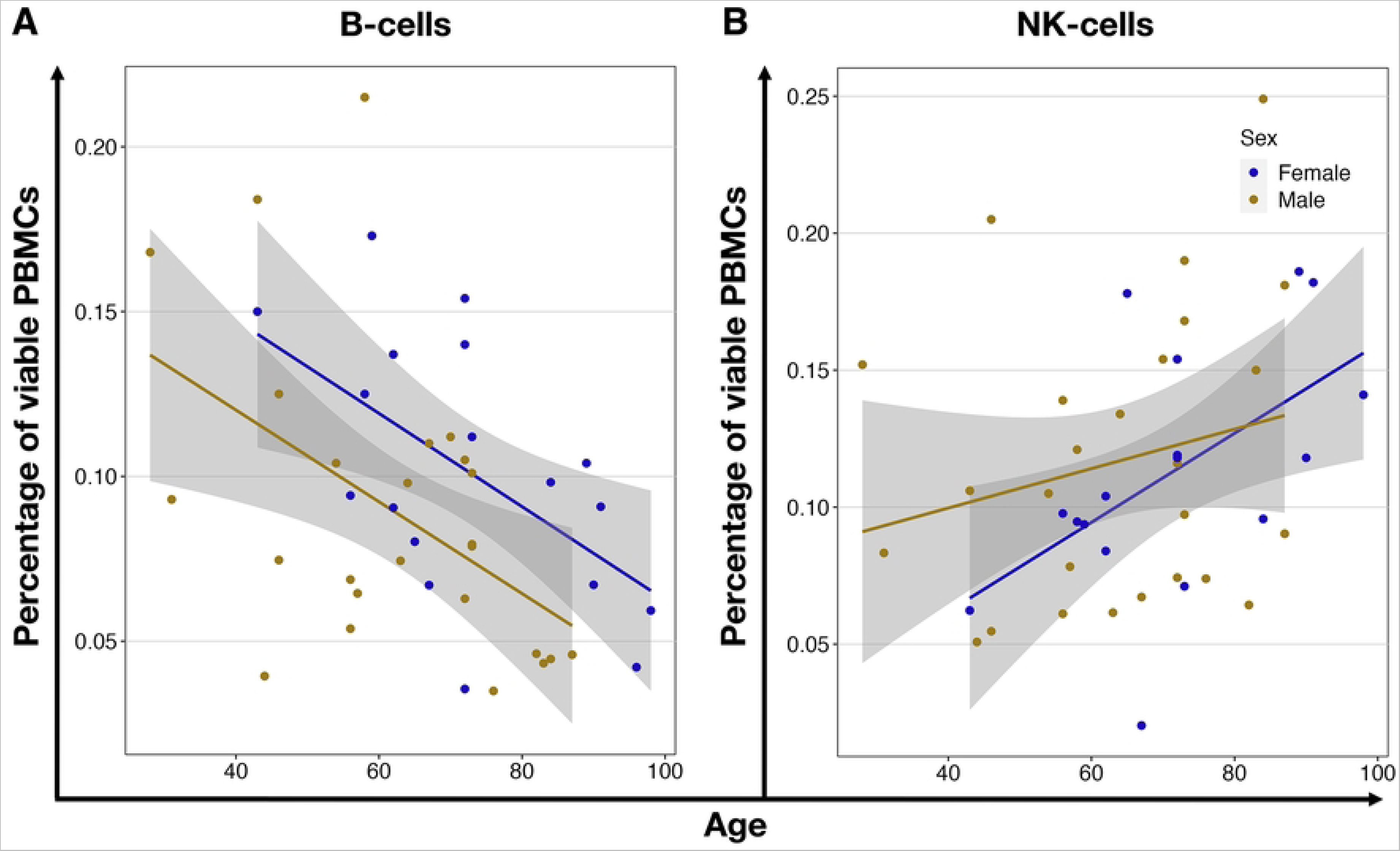
Age-related correlation of PBMCs ratios in comparison between male and female. (**A**) shows the increased (r=0.54) ratio of NK-cells and (**B**) decreased B-cells within viable PBMCs of female (r=−0.56) and male (r=−0.49) donors. Each circle represents an individual volunteer, yellow for males and blue for females. The lines show the linear regression for each biological sex, and the grey area is the 95% confidence interval.

### Subtype quantification of peripheral blood monocytes

The FACS analysis revealed that the vast majority of monocytes were represented by classical monocytes (CD14+/CD16−) (Fig. 2 C). They comprised 88.3±4.0 % and 86.1±6.2 % of all live monocytes in Adult and Older groups, respectively. Intermediate (CD14+/CD16+) monocytes were 3.6±2.1 % and 4.1±1.9 % in the Adult and Older groups, and the non-classical (CD14−/CD16+) monocytes were 8.1±2.9 % and 9.8±5.3 % in the Adult and Older groups, correspondingly (S1 Table 5). None of the monocyte subtypes differed between age groups (Supplementary Figure 1). The separate analysis, when groups were divided into males and females or correlated to age, did not reveal any age- or sex-dependent differences in the composition of monocytes either (Supplementary Table 5).

### Age-dependent expression of neuroinflammatory- and -regeneration-related genes in monocytes and their subtypes

Using the chip-based multiplex qPCR platform Fluidigm for the analysis of the expression of selected genes in monocytes, we revealed the anti-inflammatory gene, *ANXA1* (P = 0.012), and the pro-inflammatory scavenger receptor gene *CD36* (P = 0.042) which were significantly correlated to aging and both upregulated. While the upregulation of the *ANXA1* gene was also observed in the intermediate (P = 0.026) and non-classical (P = 0.004) subtypes of the monocytes (Fig 5 and Table 1), the age-related upregulation of *CD36* was not observed in any of the monocyte subpopulations (Table 1, Supplementary Table 8).

**Fig 5.**
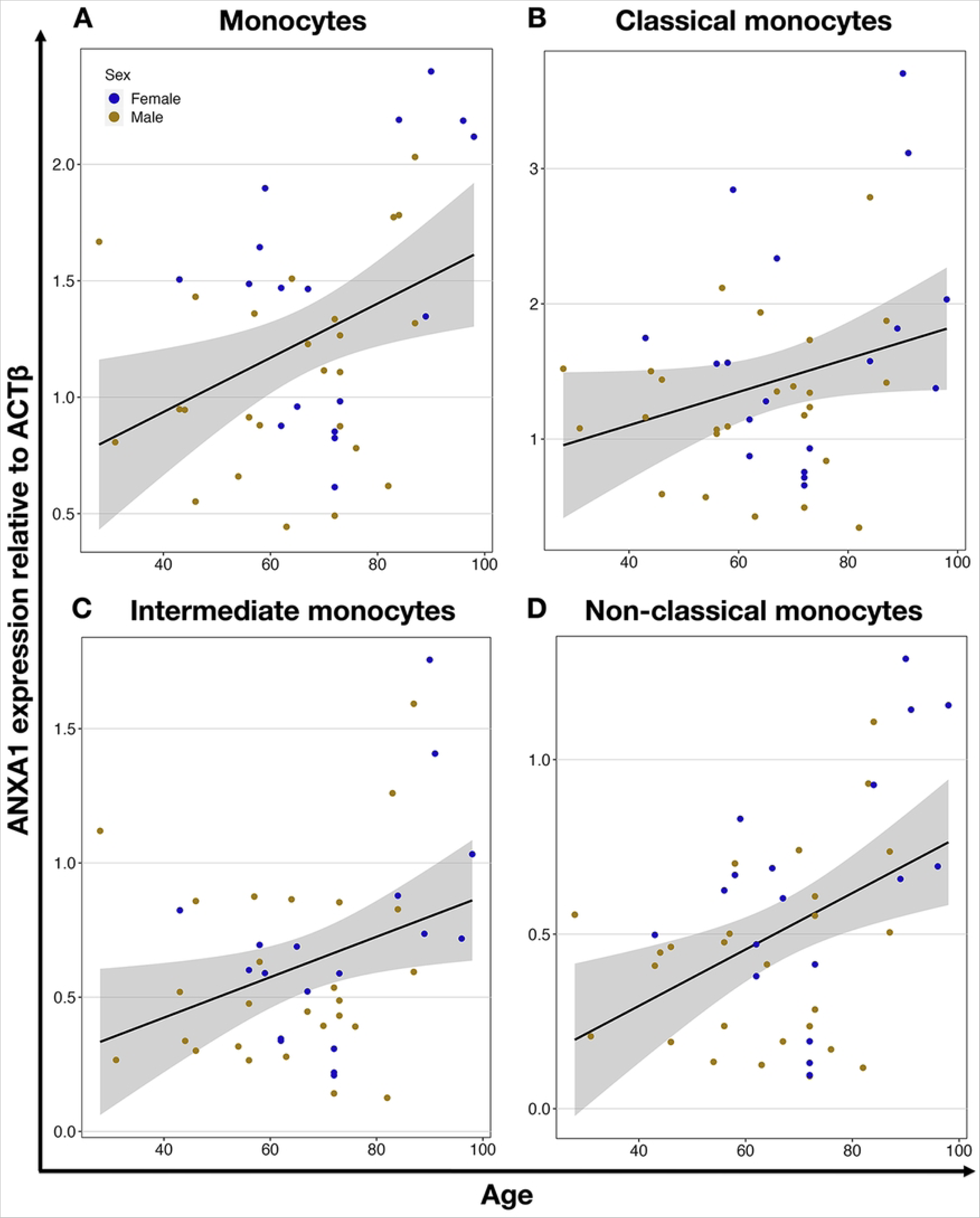
Age-dependent expression of *ANXA1* gene in monocytes and their subtypes. *ANXA1* expression in all monocytes (r=0.38) (**A**), classical (r=0.28) (**B**), intermediate (r=0.33) (**C**), and nonclassical (r=0.43) (**D**) monocytes. Each circle represents an individual donor, yellow for males and blue for females. The black line shows linear regression, and the grey area is the 95% confidence interval.

**Fig 6.**
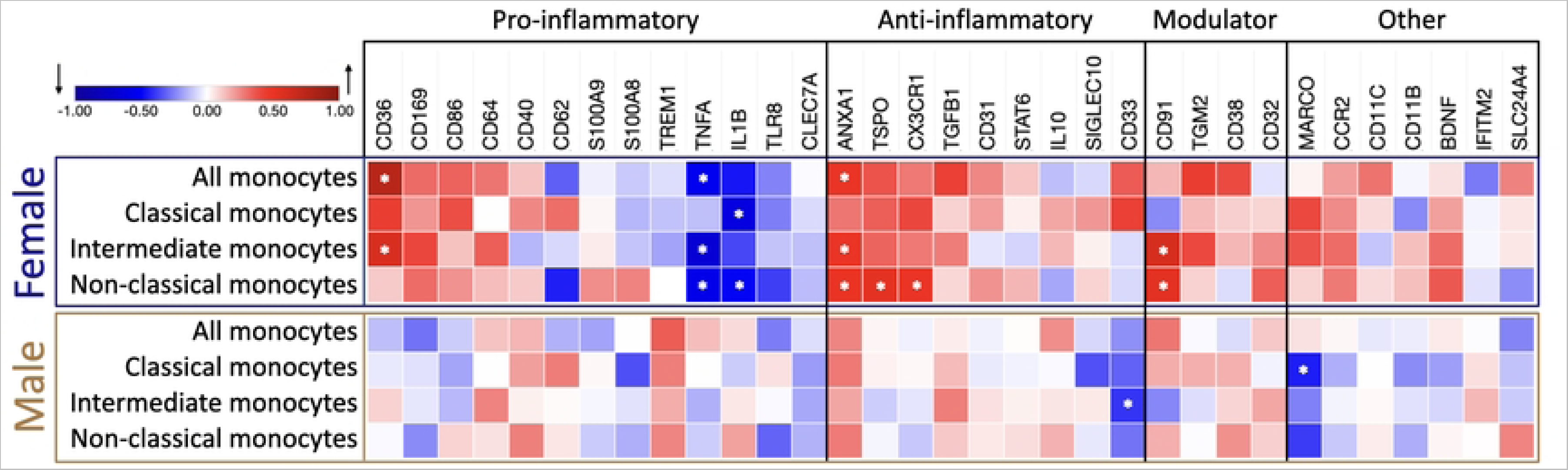
Heatmap of the relationship between age and gene expression in the different monocyte subtypes, separated by biological sex. The strength of the association of gene expression with age is represented by the Pearson correlation coefficient (r), ranging from −1, a perfect, negative correlation (blue); 0, no correlation; and 1, a perfect, positive correlation (red) of gene expression relative to *ACTβ* expression with increasing age. Genes are grouped by their most prevalent inflammatory function. Modulators have been linked to both pro- and anti-inflammatory functions. Female correlations are framed in blue (N=18), and male correlations are in gold (N=26). Significant genes, defined by a p-value below 0.05, are marked with an asterisk (*).

**Table 1.** Differentially expressed genes associated with age.

| Monocyte subtype | Gene | Function | P value | Change | Regression coefficient | 95% confidence interval |  |
| --- | --- | --- | --- | --- | --- | --- | --- |
| All | *CD36* | Pro-inflammatory, scavenger receptor | 0.042 | ↑ | 0.0019 | 0.0001 | 0.0038 |
| All | *ANXA1* | Anti-inflammatory | 0.012 | ↑ | 0.0116 | 0.0027 | 0.0206 |
| Classical | *S100A8* | Pro-inflammatory | 0.022 | ↓ | - 0.0317 | - 0.0587 | - 0.0047 |
| Intermediate | *TNFα* | Pro-inflammatory | 0.014 | ↓ | - 0.0001 | - 0.0002 | - 0.0000 |
| Intermediate | *ANXA1* | Anti-inflammatory | 0.026 | ↑ | 0.0075 | 0.0009 | 0.0141 |
| Intermediate | *TGFβ1* | Anti-inflammatory | 0.033 | ↑ | 0.0163 | 0.0014 | 0.0312 |
| Non-classical | *ANXA1* | Anti-inflammatory | 0.004 | ↑ | 0.0081 | 0.0028 | 0.0134 |
| Non-classical | *CD91* | Immune modulator | 0.031 | ↑ | 0.0001 | 0.0000 | 0.0002 |

All genes showing significant changes in the expression level in primary human monocytes and their subtypes associated with age by Pearson correlation (P-value < 0.05). An upward arrow (↑) indicates increased gene expression, while a downward arrow (↓) indicates decreased gene expression. The regression coefficient and its corresponding 95% confidence interval quantify the relative increase in gene expression with each year of age.

### Age-dependent expression of other genes in different monocyte subtypes

We further analyzed the expression of selected monocyte-related genes in the three monocyte subtypes. Interestingly, even though there were no links in the expression of the chosen genes and the age of the donors when monocytes overall were studied, we detected several genes that were strongly correlated with the donors’ age but only in defined subtypes. Namely, in the classical monocyte subpopulation, we found the pro-inflammatory gene *S100A8* (P = 0.022) to be downregulated with increasing age. In the intermediate monocyte subpopulation, we additionally found that the anti-inflammatory gene *TGFβ* (P = 0.033) was upregulated and the pro-inflammatory gene *TNFα* (P = 0.014) downregulated with increasing age. In the non-classical monocytes, we identified only one other gene: the immune modulator *CD91* (P = 0.031), which was upregulated with aging (Table 1, Supplementary Table 8).

### Sex-dependent gene expression correlates with aging

After revealing the age-dependent correlation in the expression of selected genes in monocyte subtypes, we further explored whether sex could be a factor for age-related changes (Fig. 5).

Notably, most sex-driven changes in expression of the selected genes correlating with age were found in samples from female donors. Within the female population, age significantly impacted gene expression in overall monocytes. The anti-inflammatory gene *ANXA1* (P = 0.039) and pro-inflammatory *CD36* (P = 0.0003) were upregulated with age, while the pro-inflammatory gene *TNFα* (P = 0.024) was downregulated.

When examining female monocyte subtypes, we observed that in classical monocytes, only the pro-inflammatory genes’ IL-1β (P = 0.043) and TNF*α* (P = 0.035) expression were altered with aging, displaying a significant decrease, suggesting a reduction in inflammatory activity. In the intermediate subtype of female donors, four pro-inflammatory genes had significantly changed expression with age: *TNFα* (P = 0.002), *IL-1β* (P = 0.034), and *TLR8* (P = 0.037) were downregulated, and scavenger protein *CD36* (P = 0.01) was upregulated. The anti-inflammatory *ANXA1* (P = 0.038) and the immune modulator CD91 (P = 0.007) were upregulated. In female non-classical monocytes, among the six genes that exhibited significant differential expression, all anti-inflammatory genes *ANXA1* (P = 0.035), *CX3CR1* (P = 0.0001), and *TSPO* (P = 0.024) showed upregulation, while the pro-inflammatory genes *IL-1β* (P = 0.038) and *TNFα* (P = 0.015) were downregulated with age. The immune modulator *CD91* (P = 0.023) is also upregulated in this subtype. These findings demonstrate a clear trend of increased expression in the selected anti-inflammatory genes in female non-classical monocytes.

In monocytes from male samples, only two genes in the monocytes subtype analysis were revealed as significantly downregulated: in the classical subtype, the scavenger receptor *MARCO* (P = 0.024), involved in phagocytotic activity, and in the immediate subtype, the anti-inflammatory gene *CD33* (P = 0.044) (Table 2 and Fig 5, Supplementary Table 8).

**Table 2.** Differentially expressed genes are associated with age and sex.

| Monocyte subtype | Sex | Gene | Function | P value | Change | Regression coefficient | 95% confidence interval |  |
| --- | --- | --- | --- | --- | --- | --- | --- | --- |
| All | Female | *TNFα* | Pro-inflammatory | 0.024 | ↓ | - 0.0003 | - 0.0001 | 0.0000 |
| All | Female | *CD36* | Pro-inflammatory, scavenger receptor | 0.0003 | ↑ | 0.0060 | 0.0032 | 0.0087 |
| All | Female | *ANXA1* | Anti-inflammatory | 0.039 | ↑ | 0.0246 | 0.0014 | 0.0478 |
| Classical | Female | *TNFα* | Pro-inflammatory | 0.035 | ↓ | - 0.0001 | - 0.0001 | 0.0000 |
| Classical | Female | *IL-1β* | Pro-inflammatory | 0.043 | ↓ | - 0.0022 | - 0.0044 | - 0.0001 |
| Classical | Male | *MARCO* | Phagocytotic activity, scavenger receptor | 0.024 | ↓ | - 0.0001 | - 0.0002 | - 0.0000 |
| Intermediate | Female | *TNFα* | Pro-inflammatory | 0.002 | ↓ | - 0.0003 | - 0.0004 | - 0.0001 |
| Intermediate | Female | *CD36* | Pro-inflammatory, scavenger receptor | 0.010 | ↑ | 0.0027 | 0.0007 | 0.0046 |
| Intermediate | Female | *IL-1β* | Pro-inflammatory | 0.034 | ↓ | - 0.0014 | - 0.0027 | - 0.0001 |
| Intermediate | Female | *TLR8* | Pro-inflammatory | 0.037 | ↓ | - 0.0002 | - 0.0005 | - 0.0000 |
| Intermediate | Female | *ANXA1* | Anti-inflammatory | 0.038 | ↑ | 0.0127 | 0.0008 | 0.0246 |
| Intermediate | Male | *CD33* | Anti-inflammatory | 0.044 | ↓ | - 0.0010 | - 0.0019 | - 0.0000 |
| Intermediate | Female | *CD91* | Immune modulator | 0.007 | ↑ | 0.0006 | 0.0002 | 0.0009 |
| Non-classical | Female | *TNFα* | Pro-inflammatory | 0.015 | ↓ | - 0.0002 | - 0.0003 | - 0.0000 |
| Non-classical | Female | *IL-1β* | Pro-inflammatory | 0.038 | ↓ | - 0.0004 | - 0.0007 | - 0.0000 |
| Non-classical | Female | *CX3CR1* | Anti-inflammatory | 0.0001 | ↑ | 0.0210 | 0.0130 | 0.0290 |
| Non-classical | Female | *TSPO* | Anti-inflammatory | 0.024 | ↑ | 0.0009 | 0,0001 | 0.0017 |
| Non-classical | Female | *ANXA1* | Anti-inflammatory | 0.035 | ↑ | 0.00110 | 0.0009 | 0.0211 |
| Non-classical | Female | *CD91* | Immune modulator | 0.023 | ↑ | 0.0002 | 0,0000 | 0.0003 |

All genes showing significant changes in the expression level in primary human monocytes and their subtypes separated by sex in association with age using Pearson correlation (P-value < 0.05). An upward arrow (↑) indicates increased gene expression, while a downward arrow (↓) indicates decreased gene expression. The regression coefficient and its corresponding 95% confidence interval quantify the relative change in gene expression with each year of age.

## Discussion

Our study examined the age-dependent changes in the expression of specific genes linked to inflammation and aging within peripheral blood monocyte subtypes. It aimed to explore the potential influence of sex on the observed alterations. In the pool of selected genes, we identified distinct expression patterns in male and female donors, highlighting the multifaceted nature of monocyte responses to aging, especially in females.

In-depth flow-cytometric analyses showed that the proportions of B cells, T cells, and various monocyte subtypes are stable across age groups, with classical monocytes, as expected, being the predominant population as described before [15], [19]. In agreement with prior research, our investigation revealed age-related declines in peripheral blood mononuclear cells (PBMCs) and B cells, along with an augmentation in NK-cell populations (D. Frasca et al., 2011; S. S. Gounder, B et al., 2018). Furthermore, our findings align with existing literature demonstrating an increase in the proportion of NK-cells associated with aging [47]. When we investigated the donors separated by biological sex, this age-associated rise in NK-cell proportions reached statistical significance solely in females, substantiating earlier observations [48]. In the sex-stratified analysis, B-cell counts also significantly decreased in both, females and males with age, consistent with previous studies [49].

When we separated the donors into two groups based on age (under and above 65 years), there were no significant differences in the ratio of different blood cells and monocyte subtypes, as well as the expression of several neuroinflammation and regeneration-related genes between these groups. However, the correlation analysis indicates an age-related change in the ratio of cells and gene expression with aging when considering age as a continuous variable. This suggests that while there may not be a strong difference between specific age groups, there are significant changes as individuals age continuously. This observation could indicate that each human has a specific aging curve of the immune system and monocyte subtypes expressing neuroinflammation and regeneration-related genes. Introducing arbitrary cuts in the age for separating adult and old populations might be biased and hinder the accurate picture based on individual variability. Therefore, correlation analysis of age-related changes holds more potential than separating data based on arbitrarily introducing an age cut-off point. This approach could help to reveal valuable data for understanding the dynamics of biological changes in aging individuals.

For the first time, we demonstrated increased expression of the anti-inflammatory *ANXA1* gene in the CD91+ population of monocytes, driven by higher levels of this gene in intermediate and non-classical subtypes of monocytes in relation to aging. *ANXA1* is a gene coding for Annexin A1, which belongs to the annexin superfamily of calcium- and phospholipid-binding proteins [50]. The Annexin A1 protein exerts anti-inflammatory effects through the G-coupled formyl peptide receptor type-2 with a phospholipase A2 inhibitory activity (reviewed by Perretti and D’Acquisto [51]).

While the involvement of *ANXA1* in regulating inflammation is well-documented, the relevance of *ANXA1* expression in longevity and healthy aging is unknown, and our finding implicates its potential connection with the aging process. Notably, *ANXA1* has been demonstrated to be involved in a variety of biological processes, such as regulation of macrophage phagocytosis and neutrophil migration, acute [52] and chronic [53] inflammation, and ischemia/reperfusion injuries [52, 54, 55]. The restoration of plasma *ANXA1* levels after stroke is indicative of a favorable recovery in stroke patients, suggesting its potential as a biomarker and valuable prognostic tool [56]. It has been shown that the classical monocyte subtype conveys detrimental effects after stroke, including stronger interaction with platelets [34]. In contrast, non-classical and intermediate monocytes are beneficial with a phenotype that could promote tissue repair and angiogenesis. The negative outcome after stroke is increased with aging [57], and the lack of the increase of ANEXIN A1 expression in classical monocytes might be a confounding factor.

We detected increased expression of the pro-inflammatory *CD36* gene in all monocytes with aging. *CD36*, a multi-ligand scavenger receptor expressed across diverse cell types, operates context-dependently, manifesting a robust pro-inflammatory response when expressed in monocytes and macrophages. *CD36* plays a role in various biological functions. It mediates innate immunity, participating in the assembly of inflammatory pathways and contributing to reactive oxygen species (ROS) production [58], and plays a role in macrophage phagocytosis during the resolution phase of ischemic stroke in mice [59].

Inflammation is vital in maintaining the body’s homeostasis and promoting recovery after injury. However, if inflammation becomes excessively aggressive or persists without being resolved, it can result in profound tissue damage [60]. The observed upregulation of the *CD36* gene in all monocytes suggests a potential shift towards a more pro-inflammatory phenotype during aging. In contrast, the concurrent upregulation of the *ANXA1* gene expression suggests an anti-inflammatory tendency. Therefore, it is plausible that the increased expression of *ANXA1* and *CD36* may act in concert, potentially offsetting each other and pointing towards dysregulation in inflammation mechanisms during aging.

In the classical monocytes, but not in other subtypes, we detected the downregulation of the *S100A8* gene, a calcium-binding protein belonging to the S100 family [61]. The S100a8 (calgranulin A) and S100a9 (calgranulin B) proteins are constitutively expressed in neutrophils and monocytes. They are potential biomarkers for inflammation-associated diseases and key inflammatory regulators with the capacity to initiate and react to signals associated with inflammation [62]. In contrast with our findings, it has been shown that the increase in expression of *S100A8/A9*, particularly *S100A9*, represents a characteristic of aging across various mammalian tissues [63]. This phenomenon involves diverse cell types, including those in the blood and the central nervous system. In healthy human donors, decreased levels of the *S100A8/S100A9* were found in the serum of the elderly compared to the young individuals. However, data regarding the expression of *S100A8* in monocytes with aging are not entirely consistent across studies. It has been shown that expression of *S100A8* in classical and intermediate monocytes individually is higher compared to the other two subtypes but in non-classical is lower [64]. Moreover, *S100A8* also differentially responds to the activation of monocyte subtypes though with similar changes in cells from young and old donors [64].

Nevertheless, it is important to note that such discrepancy could be attributed to the fact that these studies did not specifically focus on monocytes; instead, they analyzed whole blood or other tissues. Currently, aging is often associated with increased pro-inflammatory cytokines in blood plasma. However, our data underscores the importance of discerning the distinct contributions of various monocyte classes to the aging phenomenon.

In the intermediate monocytes, in addition to the *ANXA1*, the anti-inflammatory *TGFβ1* gene was upregulated, and the pro-inflammatory gene TNF-α was downregulated with increasing age. Although intermediate monocytes act as antigen-presenting cells, secrete cytokines, and regulate apoptosis, their precise role in immunity appears elusive [65]. The concomitant age-dependent anti- and pro-inflammatory gene expression changes suggest a potential contribution to immune dysregulation and inflammation of the intermediate monocytes.

In non-classical monocytes, only two genes, *ANXA1* and *CD91*, were upregulated with aging. In the present study, we have successfully used *CD91*, the adhesion molecule, for the identification of human monocytes in a more accurate manner instead of solely relying on the *CD14* and *CD16* expression [66]. *CD91* has been identified as a receptor in antigen-presenting cells, including monocytes, which plays a crucial role in the innate and adaptive immune response [67]. In contrast with recently published literature showing a higher expression of *CD91* in classical and intermediate monocytes through flow-cytometry analysis as a predictor of age advancement, we found that non-classical monocytes express the highest levels of *CD91* [68]. However, it should be emphasized that the differential techniques utilized in our study may have contributed to the observed disparities, emphasizing the importance of methodological considerations in interpreting and contextualizing research outcomes.

After revealing the age-dependent correlation of gene expression in monocyte subtypes, we further explored whether biological sex could contribute to the observed age-related changes. Classical monocytes isolated from male donors show downregulation of *MARCO*, the macrophage receptor with collagenous structure, and intermediate monocytes downregulated the anti-inflammatory marker *CD33*. Notably, the female donors display changes in the phenotype of all three monocyte subtypes with age. At the same time, the classical monocytes showed downregulation, specifically of pro-inflammatory genes *TNF-α* and *IL-1β*, and the intermediate and non-classical subclasses exhibited significant alterations in the expression of multiple genes, indicating the acquisition of an ambiguous pro- and anti-inflammatory phenotype with aging. Importantly, we report for the first time a significant downregulation of the *TLR8* gene expression and an upregulation of the *TSPO* gene expression in intermediate monocytes and non-classical monocytes, respectively, providing valuable insights into the dynamic changes occurring at the molecular level within specific monocyte subtypes during the aging process.

*TLR7* and *TLR8* are crucial components of the innate immune response, recognizing RNA degradation products from pathogens. *TLR8*, found in monocytes and dendritic cells, is not subject to X chromosome inactivation in certain cells, possibly leading to higher *TLR8* levels in females. This has implications for antiviral and antibacterial responses, as well as susceptibility to inflammatory and autoimmune diseases, highlighting *TLR8*’s role in immune regulation and disease vulnerability [69]. In a recent study, TLR8 protein expression was reported to be higher in all subsets of female monocytes compared to their male counterparts in healthy blood donors aged 16-44 years, and flow cytometry analysis revealed that, on average, 60% of non-classical monocytes positively stained for TLR8 [70]. Intriguingly, this suggests a potential alteration in pathogen response and increased susceptibility to autoimmune and inflammatory diseases in aging individuals.

We found that *TSPO* gene expression is upregulated in female donors with age, exclusively in the non-classical monocytes. The mitochondrial membrane protein, translocator protein (TSPO), is found on both the mitochondrial and plasma membranes of diverse cell types, including brain microglia cells and circulating lymphocytes and monocytes [71]. Previous *ex-vivo* human studies demonstrated the impact of various TSPO ligands on the monocyte chemotaxis [72].

Recently, it has been hypothesized that peripheral monocytes’ recruitment through the blood-brain barrier may contribute to the increase of TSPO levels in the central nervous system observed in the context of inflammation and Alzheimer’s disease [73]. Our findings reveal a novel association between age, *TSPO* gene expression, and the unique functional characteristics of non-classical monocytes. The upregulation of *TSPO* in female donors could play a role in highly specialized functions of non-classical monocytes, which include trans-endothelial migration, phagocytosis, and viral response, and may contribute to the overall immune surveillance in aging individuals.

The newly coined term ”inflammaging” indicates the intricate relationship between inflammation and aging, wherein chronic low-grade inflammation accelerates the aging process while aging itself fuels a pro-inflammatory environment. This phenomenon is proposed to be driven by various mechanisms inducing pro-inflammatory cytokine secretion, cellular senescence of immune and non-immune cells, increased release of inflammatory mediators via a senescence-associated secretory phenotype, and perturbed gut integrity facilitating bacterial product entry into the circulation [74]. However, a persistent state of inflammation does not affect elderly individuals in the same fashion, and healthy individuals who have reached the remarkable age of 100 years appear to effectively compensate for such inflammatory responses [75].

Aging exhibits gender specificity, and in women, it is characterized by the occurrence of menopause, typically taking place between 45 and 55 years of age globally (World Health Organization, 2022). Notably, our study shows that female donors display robust age-related changes in the phenotypes of all three monocyte subtypes. Our study’s age range of female donors extends from 43 to 98 years, suggesting a potential overrepresentation of peri-menopausal and menopausal women within the female sample group. Previous studies have shown disruptions in the cyclic pattern of circulating estrogen, a potent anti-inflammatory agent, [76] during the menopausal shift. This activates broader innate and adaptive immune reactions in the body [77], elevating levels of chronic systemic inflammation [78], increasing the risk of cardiovascular diseases [79] and making the brain more susceptible to ischemic damage. This highlights the potential complex interplay of monocyte subtypes in the context of aging.

In a previously published study exclusively examining females comparing pre-menopausal (50.0 ± 3.1 years old) and post-menopausal (52.0 ± 1.7 years old) women, it was noted that menopause influences the functional state of circulating monocytes [80]. Our study extends this knowledge, demonstrating that females exhibit more pronounced changes in gene expression among monocyte subtypes than males, emphasizing the significant impact on females in the context of aging. Consequently, comprehending the biological mechanisms underlying the transition to menopause can contribute to the development of strategies aimed at protecting women from health complications associated with menopause.

In conclusion, our data show the regulation of selected inflammation- and regeneration-related genes in human monocyte subtypes in the context of age and sex. This highlights the importance of considering the individual characteristics of patients with inflammation-related diseases, including sex and age. Moreover, the monocyte-related inflammatory response should probably be considered from a monocyte subtype perspective. However, future studies must fully reveal the functional and pathological significances of all differential gene expressions observed in the present study. Our future research aims to expand beyond healthy volunteers, focusing on comprehensive gene studies in ischemic stroke patients. Such investigations have the potential to unveil a broader spectrum of genomic changes associated with the severity and prognosis of ischemic stroke, with particular consideration given to the age and sex of the patients.

## Acknowledgments

This work was made possible by the contribution of doctors and research nurses at Skåne University Hospital and the support of the Lund Stroke Register (LSR). We thank the participants for donating blood samples and contributing to this study. The authors thank the Lund Stem Cell Center FACS and Bioinformatics Facilities for their technical support and input. We thank Prof. Dennis Sandris Nielsen and Dr. Bashir Aideh at the Department of Food Science at the University of Copenhagen for providing access to the Fluidigm system.

